# Learning-Based Estimation of Fitness Landscape Ruggedness for Directed Evolution

**DOI:** 10.1101/2024.02.28.582468

**Authors:** Sebastian Towers, Jessica James, Harrison Steel, Idris Kempf

**Affiliations:** Department of Engineering Science, University of Oxford, Parks Road, Oxford OX1 3PJ, UK

**Keywords:** Protein engineering, directed evolution, machine learning

## Abstract

Directed evolution is a method for engineering biological systems or components, such as proteins, wherein desired traits are optimised through iterative rounds of mutagenesis and selection of fit variants. The process of protein directed evolution can be envisaged as navigation over high-dimensional landscapes with numerous local maxima, mapping every possible variant of a protein to its fitness. The performance of any strategy in navigating such a landscape is dependent on several parameters, including its ruggedness. However, this information is generally unavailable at the outset of an experiment, and cannot be computed using analytical methods. Here we propose a learning-based method for estimating landscape ruggedness from a mutating population, using only population average performance data. This method uses a short period of exploration at the beginning of an experiment to predict the ruggedness, subsequently guiding the choice of high-performing parameters for directed evolution control. We then simulate this approach on two real-world protein fitness landscapes, demonstrating an improvement upon the performance of standard strategies, particularly on rugged landscapes. In addition to improving the overall outcomes of directed evolution, this method has the advantage of being readily deployable in laboratory settings, even in configurations that exclusively capture average population measures. Given the rapidly expanding application space of engineered proteins, the products of improved directed evolution are relevant in medicine, agriculture and manufacturing.

## 1. Introduction

Proteins are fundamental biological components that perform various functions within living organisms. Proteins are variable-length chains of sub-units known as amino acids, for which there are 20 different types. The length and sequence of amino acids in each protein is dictated by a gene, which is composed of DNA. In an ideal world, new proteins could be engineered by *de novo* protein design (Jumper et al., 2021; Baek et al., 2021), which aims at understanding how the sequence of amino acids maps to structure and function of the protein. However, due to the huge combinatorial possibilities (20^*N*^ for a protein of length *N* ), *de novo* protein design remains difficult, even with recent advances in computing power and deep learning methods (Dauparas et al., 2022; Ferruz et al., 2022; Singer et al., 2022; Anishchenko et al., 2021; Wicky et al., 2022).

In contrast, directed evolution is a process by which biological components, such as proteins, are engineered and improved through iterative rounds of selection and mutagenesis, emulating the natural evolution process (Arnold, 1998). Directed evolution has had many successful applications, including the development of drugs (Nixon et al., 2014) and biofuels (Heater et al., 2019). The problem of directed evolution can be interpreted as navigation over a protein *fitness landscape* (Wright, 1932). Fitness landscapes are high-dimensional structures in which the sequence of the protein represents a coordinate that maps to a fitness value, which, in the context of this work, is defined as the property the directed evolutionary process aims to optimise. Fitness landscapes are known to exhibit variable degrees of ruggedness, which can create local optima that constrain paths of evolution (Wu et al., 2016).

The standard approach to directed evolution is to select the best performing variants with each iteration. However, this approach can be prone to getting trapped in local optima (Carpenter et al., 2023). In recent years, numerous optimisation methods have been developed, leveraging machine learning to actively navigate a protein fitness landscape (Wu et al., 2019; Wittmann et al., 2021; Yang et al., 2019; Frisby and Langmead, 2021; Hu et al., 2023; Fox et al., 2003). Although effective, these optimisation approaches require sequencing of the entire population of variants with each iteration. This makes them labour- and resource-intensive, and not applicable to novel continuous directed evolution methods where DNA sequence data is not generally available during an experiment (Molina et al., 2022). In a previous work, we proposed strategies that can be used to optimise directed evolution without the need to sequence all variants (James et al., in press). We found, however, that the performance of each strategy is dependent on the properties of the underlying landscape, information that is generally not available at the outset of an experiment.

It has previously been shown that neural networks can be used to infer evolutionary parameters, such as rate of accumulation of beneficial mutations (Avecilla et al., 2022). Statistical models have also been generated for inferring protein fitness landscape properties from directed evolution trajectories with sequencing information (D’Costa et al., 2023). In this paper, we use a neural network to estimate properties of a protein fitness landscape without sequencing information. The method requires measurements of the average fitness from a population mutating away from a starting point, which is easily implementable in various experimental configurations. Such data can be collected by mutating a single population of bacteria (e.g. by UV), and recording the average population fitness value (e.g. fluorescence) at regular intervals. The resulting measurements are input into a fully connected neural network (FCN), which has been pre-trained on theoretical fitness landscapes to predict ruggedness. The use of an FCN is required as fitness landscapes represent a highly non-linear mapping from DNA sequence to protein functions and properties, which cannot be captured by traditional methods such as linear regression. This estimate is then used to select directed evolution control parameters that are sensitive to the ruggedness, such as identified in our previous research (James et al., in press). We analyse the prediction accuracy of the FCN with respect to the number of fitness values provided, and demonstrate that the ruggedness can be reliably estimated, resulting in a performance increase of directed evolution experiments in highly rugged landscapes. Finally, we apply our estimation procedure to two real-world landscapes and demonstrate that it can be used in practical settings, even though trained on theoretical models.

The paper is organized as follows. Section 2 frames directed evolution as a control problem. In Section 3, the proposed method for parameter inference is developed. The paper is concluded by evaluating the proposed method on empirical landscapes.

## 2. Problem Formulation

Directed evolution can be represented by the feedback loop shown in Fig. 1a, which includes measurement noise. Let 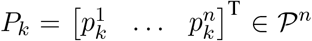 denote the *n* members of the evolved population at iteration *k* ∈ ℕ, *F* : 𝒫 ↦ ℝ the fitness function, *S* : ℝ^*n*^ ↦ {0, 1{^*n*×*n*^ the selection process, and *M* : 𝒫 ↦ 𝒫 the mutagenesis process. For a gene of *N* loci, each with *A* possible alleles, P represents a discrete sequence space with *A*^*N*^ different sequences, *A* ∈ ℕ^+^. The function *F* measures the performance of a population member and has multiple local optima in general. The selection process takes *F* (*P*_*k*_), where *F* is applied element-wise, as inputs, and outputs a selection matrix *S*(·) with exactly one 1 per row, so that the selected variants are obtained from *S*(*F* (*P*_*k*_))*P*_*k*_. Note that the sequences are not observed directly in practice, only their fitnesses *F* (*P*_*k*_). The function *M* is modelled as a stochastic function that changes each element of 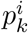 with probability *θ* and leaves it unchanged with probability 1−*θ*, so that the number of mutations approximately follows a Poisson distribution with mean *Nθ*.

**Figure 1.**
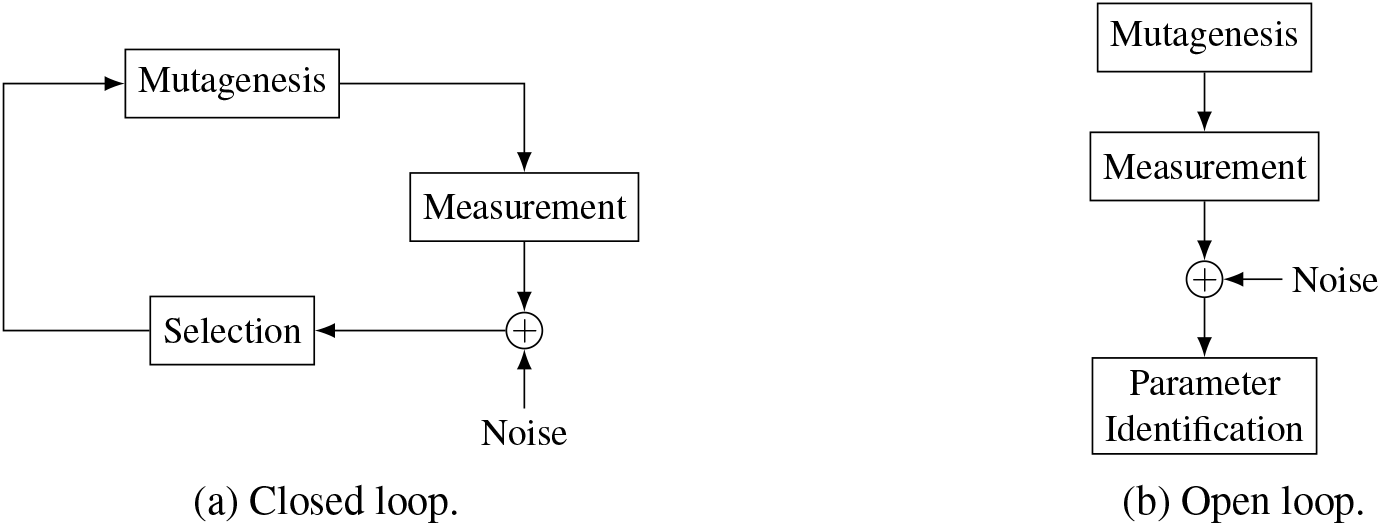
Block diagrams for directed evolution in closed-loop and open-loop configurations.

With these definitions, the process from Fig. 1 can be modelled as

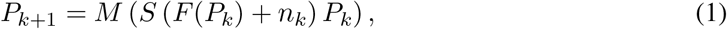

where *n*_*k*_ ∈ ℝ^*n*^ refers to the noise, and the functions *F* ( . ) and *M* ( . ) are applied element-wise. For the remainder of the paper, the effect of noise is ignored. The aim of directed evolution is to maximise the maximum fitness of the population, i.e.

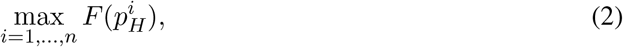

where *H* ∈ ℕ^+^ is the fixed duration of the experiment. Because *F* lacks strong structure in general, *M* is a stochastic function, and the sequences 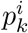 are not directly observed, problem (2) cannot be solved using standard optimisation techniques.

The standard approach to selection in directed evolution is to just choose the fittest variants in each generation. This approach is prone to getting trapped in local optima, particularly in rugged landscapes (Carpenter et al., 2023). In order to reduce this propensity, in a previous work we proposed an alternative function for selection shown in Fig. 2a (James et al., in press). The selection function is defined by two parameters: a threshold fitness percentile, *t* ∈ [0, 1], above which variants are always selected, and a base chance of selection, *b*∈ [0, 1], for variants with lower fitness percentiles. The expected fraction of cells *f* selected at each iteration is represented by the shaded area in Fig. 2a and given by *f* := 1− *t*(1−*b*). Throughout the paper it is assumed that *f* = 0.25 (James et al., in press), so that for a given *b*∈ [0, *f* ], *t* = (1− *f* )*/*(1− *b*). This selection function trades off exploration with exploitation, and is therefore less prone to getting trapped in local optima than the aforementioned standard approach. Depending on the properties of fitness landscape, it has been shown that the choice of *b* significantly affects the outcome of the experiment (James et al., in press). While low base chances (greedy selection) perform well on landscapes with few maxima, high base chances benefit the outcome of the experiment on rugged landscapes. This is captured in Fig. 2b, which shows the best-performing *b* as a function of the gene length *N* and a ruggedness measure *K/N* on the theoretical *NK* landscape (Section 3.1) for a fixed number of iterations *H* = 100. For heavily rugged landscapes (*K/N* ≈ 1), high base chances are favourable, whereas for less rugged landscapes (*K/N* ≪ 1), low base chances are favourable.

**Figure 2.**
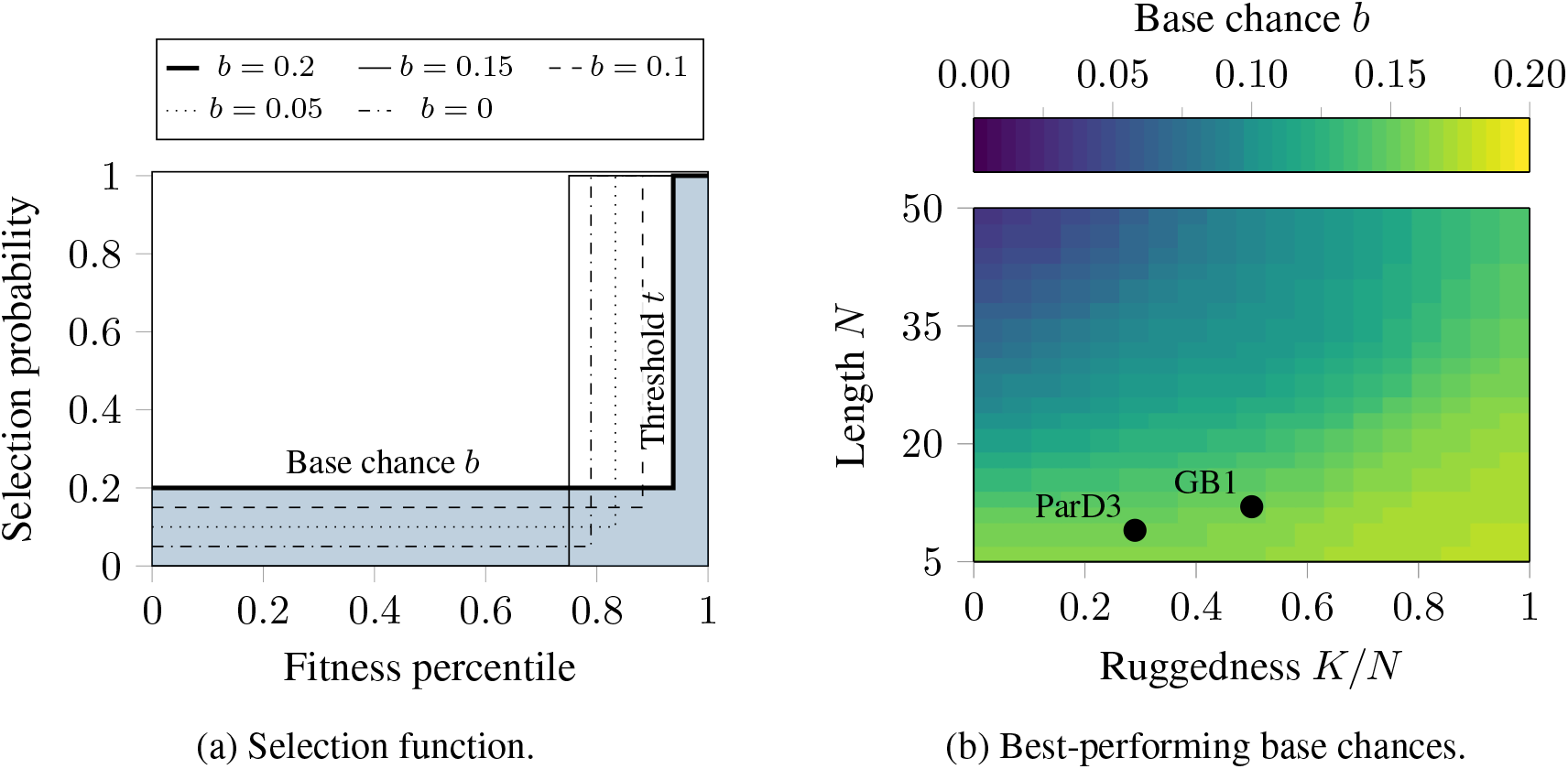
Stochastic selection functions (with *t* = (1− *f* )*/*(1− *b*) and *f* = 0.25) and corresponding best-performing base chances on the *NK* landscape (adapted from (James et al., in press)). Figure (b) also shows the estimated *K/N* for the GB1 and ParD3 landscapes using the method from Section 3.

## 3. Estimating Landscape Ruggedness

The fitness landscapes of biological entities evolved in real-world experiments are not known a priori. The question arises whether landscape properties can be inferred from fitness measurements taken before the start of the experiment in order to choose appropriate control parameters for selection and mutagenesis. To achieve this, a method for inferring a ruggedness measure of the landscape is developed, which is in turn used to select the parameter *b*. Given that *F* can be arbitrarily complex, an FCN is trained to estimate the ruggedness from fitness values measured in the open-loop configuration from Fig. 1b, where the selection procedure has been omitted. First, information on the landscape is collected by repeatedly mutating the population and measuring the average fitness. Second, these measurements are preprocessed and fed into the trained FCN, which outputs an estimate of the ruggedness. Finally, the ruggedness estimate is used to choose an appropriate base chance from the look-up table in Fig. 2b.

### 3.1. *NK* Landscapes

The successful training of the FCN hinges on the availability of data. At present, there exist few empirical data sets that map sequence space to fitness measurements (Section 4). These empirical data sets are therefore not in sufficient quantity for training the FCN, nor are they accompanied by a clear definition of ruggedness for training. This problem can be circumvented by using theoretical models of fitness landscapes, such as Fisher’s geometric model (Tenaillon, 2014; Fisher, 1931), the holey landscape model (Gavrilets, 1997), the multilinear model (Hansen and Wagner, 2001), the Rough Mount Fuji model (Neidhart et al., 2014) and the *NK* model (Kauffman and Levin, 1987; Kauffman and Weinberger, 1989), which is used here on account of its tuneable ruggedness and implementation of epistasis (interaction between sub-units, which is prevalent in a protein context).

In the *NK* model, each gene is represented by a sequence of length *N* . Every site interacts with *K* other sites in the gene, influencing the resulting fitness. The number of interactions, determined by *K*, is what allows ruggedness to be tuned. When *K* = 0, the landscape is linear and has a single peak. The other extreme, *K* = *N* −1, is maximally rugged, with all fitness values independent from one another. Values of *K* between the two interpolate between these two extremes. Note that, even though the underlying generation process is known, finding the global optimum of an *NK*-landscape is an NP-hard problem for *K >* 1 (Wright et al., 2000).

### 3.2. Training the Fully Connected Neural Network

To estimate the ruggedness, the population is mutated for *G* generations in the open-loop configuration from Fig. 1b, so that *P*_*k*+1_ = *M* (*P*_*k*_) = *M* ^(*k*+1)^(*P*_0_). To account for numerical differences between the landscapes, the measurements 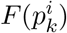 are normalised by the observed mean, *ū*, as 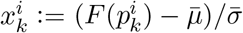, where 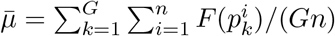 and 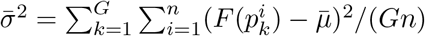.

The mean, *μ*_*k*_, of the population fitness of generation *k* are then computed as 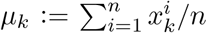 from which the features provided to the FCN are computed as

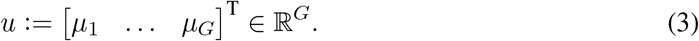

The choice (3) is designed to be as simple as possible, while capturing factors relevant to ruggedness estimation. Fig. 3a shows two example trajectories for a smooth (*K/N* = 0.2) and a rugged (*K/N* = 0.8) landscape with *N* = 100, *Nθ* = 0.5, and with *n* = 50 (solid) and *n* = 4000 (dotted). It can be seen that both trajectories converge to the overall mean fitness, 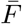, of their corresponding landscape, but at different speeds. For a rugged landscape, *μ*_*k*_ converges more quickly to 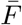 than for a smooth landscape. This is formalised for the *NK* landscape in the following proposition:

**Figure 3.**
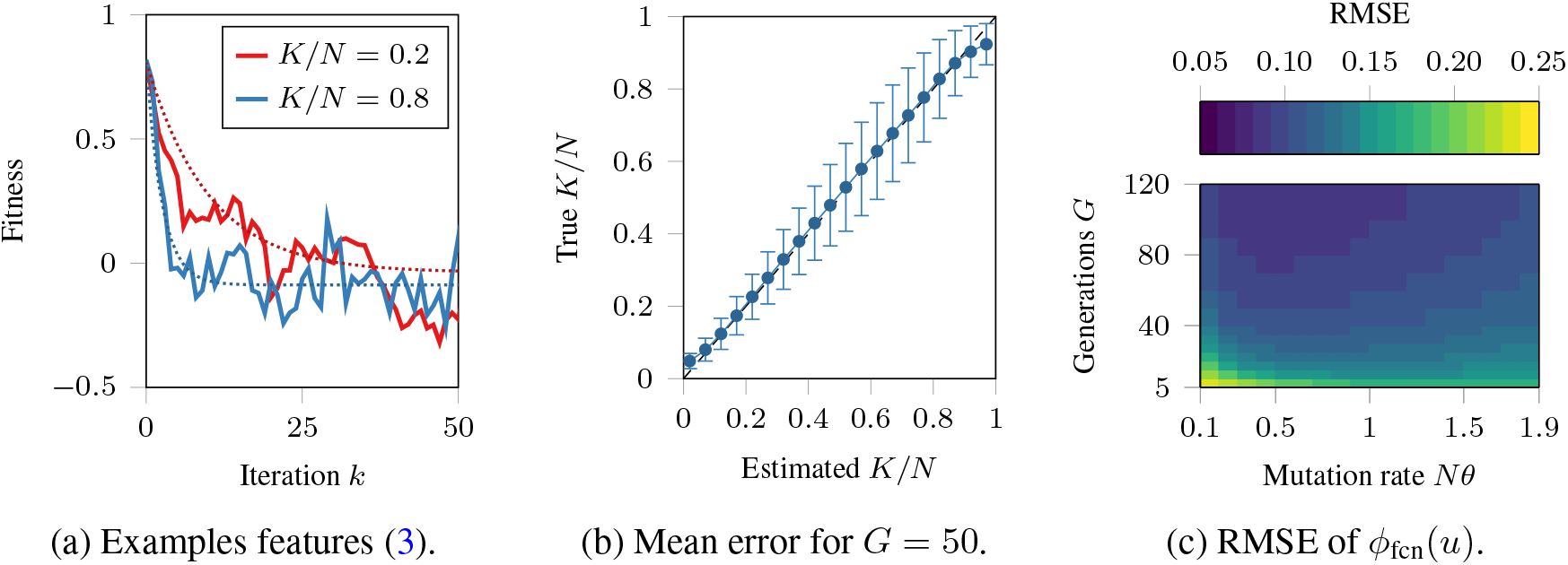
Example features (*N* = 100 and *n* = 50) and performance of the ruggedness estimator evaluated on a test set with 1.2 × 10^6^ datapoints.

#### Proposition 1

*Let F* : *A*^*N*^ → ℝ *be an NK landscape, and let* 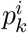 *be a single cell mutating at rate θ per generation. Then:*

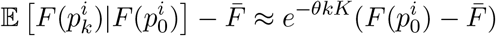

**Proof** See Appendix A.

In particular for large *n, μ*_*k*_ can be interpreted as an population estimate of 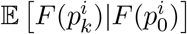, so that 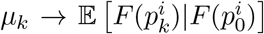 as *n* →∞. As *K* is a measure of ruggedness, this implies that the more rugged the landscape, the more quickly *μ*_*k*_ converges to 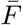. Note that the approximation from Proposition 1 coincides with the dotted line from Fig. 3a, which corresponds to (3) with *n* = 4000.

In order to infer *K/N* from (3), an FCN, *ϕ*_fcn_(*u*) ≈𝔼 [*K/N u*], is trained on a dataset of 1.2 × 10^6^ (*u, K/N* ) pairs generated from a distribution of *NK* landscapes. The FCN has two hidden layers of size 128 and is trained using gradient descent. The main computational cost is data generation, which takes 10 min using an NVIDIA A40, whereas the training of the FCN takes 2 min. The estimation error of *ϕ*_fcn_(*u*) obtained on a test set with 1.2 10^6^ datapoints is shown in Fig. 3b for different values of *K/N* and *G* = 50. It can be seen that *ϕ*_fcn_(*u*) performs well for extreme values of *K/N*, but worse for intermediary values.

The accuracy of *ϕ*_fcn_(*u*) is further analysed in Fig. 3c for 5≤ *G* ≤ 120 and mutation rates 0.1 ≤ *Nθ* ≤ 1.9, which shows the root mean square error (RMSE) averaged over the values of *K/N* marked in Fig. 3b. Fig. 3c shows that the accuracy increases as *G* does, but plateaus quickly, which is related to the convergence properties of *μ*_*k*_ shown in Fig. 3a. Fig. 3c also shows that a lower mutation rate requires a larger *G* for a high accuracy.

## 4. Translation to Empirical Landscapes

The performance of *ϕ*_fcn_(*u*) is tested on two different empirical landscapes. Empirical landscapes are experimental data sets describing the fitness of all possible variants of a protein region. Measurement of such fitness landscapes can be a highly resource-intensive task, as they increase in size exponentially with the addition of each amino acid position. At present, there is a limited number of empirical protein fitness landscapes, with the largest combinatorially complete example not exceeding four amino acid positions (Weinreich et al., 2006; Khan et al., 2011; Chou et al., 2014; Bank et al., 2016; Wu et al., 2016; Lite et al., 2020; Papkou et al., 2023). There is a necessity, therefore, to train *ϕ*_fcn_(*u*) using theoretical *NK* landscapes, and to reserve the empirical landscapes for testing. *NK* landscapes are a simplistic representation of true protein fitness landscapes, given the fact that *K* is a fixed constant and not variable over the protein, and that the distribution of fitness values is normal, as opposed to being heavily skewed towards zero. Despite this, it is found that the model fares well when applied to real fitness landscapes, and may improve with more tailored theoretical fitness landscapes.

The first empirical fitness landscape the ruggedness estimator is tested on is that of a four amino acid region of GB1 (20^4^ combinations) (Wu et al., 2016). GB1 is an antibody-binding protein isolated from Streptococcal bacteria. The fitness values of this landscape correspond to how well each GB1 variant binds to the antibody. The second empirical landscape tested on is that of a three amino acid region of an antitoxin protein known as ParD3 (20^3^ combinations) (Lite et al., 2020). Fitness values in this landscape correspond to strength of binding to the toxin ParE2.

Ruggedness (*K/N* ) estimation is performed on GB1 and ParD3 landscapes using a mutating population of size *n* = 50, mutating from the natural (wildtype) sequence over *G* = 50 generations, with a mutation rate *Nθ* = 0.5. As the process underlying the ruggedness estimation is stochastic, *ϕ*_fcn_(*u*) is evaluated 100 times. The FCN *ϕ*_fcn_(*u*) estimated *K/N* values of 0.5 0.11 and 0.28 0.04 on GB1 and ParD3, respectively (see Fig. 2b). The estimated *K/N* values are combined with *N* to look up an estimate for optimal base chance. Here, *N* corresponds to the length of the DNA sequence for the protein mutating region, so that *N* = 12 for GB1 and *N* = 9 for ParD3. The inferred base chance values are *b*_fcn_ = 0.157 on GB1, and *b*_fcn_ = 0.156 on ParD3.

Finally, directed evolution simulations are performed on GB1 and ParD3 using the selected base chance values and compared in Fig. 4. In each case, the simulation is compared to the standard approach to directed evolution, which is to select only the top fraction of variants each generation (*b* = 0), as well as to the “optimal” base chance *b*_opt_ obtained in the same way Fig. 2b was obtained. On GB1, a 5.6 % improvement in fitness after 100 generations is observed. As a landscape that is predicted to be more rugged, it is proposed that this improvement is due to the increase in *b*, reducing the propensity to get trapped in local optima. On closer inspection, this was confirmed as the mean fitness of the new strategy corresponds to between the highest and second highest peak on the landscape, whereas the mean fitness of the standard approach corresponds to between the second and third highest peak on the landscape. On ParD3, all strategies achieved the global maximum (1.023) within ∼10 generations. This supports the estimated lower *K/N* value (0.28), which suggests that ParD3 is a less rugged landscape and thus easier to navigate. In each case, the strategy that uses *b*_fcn_ either matches or out-performs the standard approach, and matches the strategy that uses *b*_opt_.

**Figure 4.**
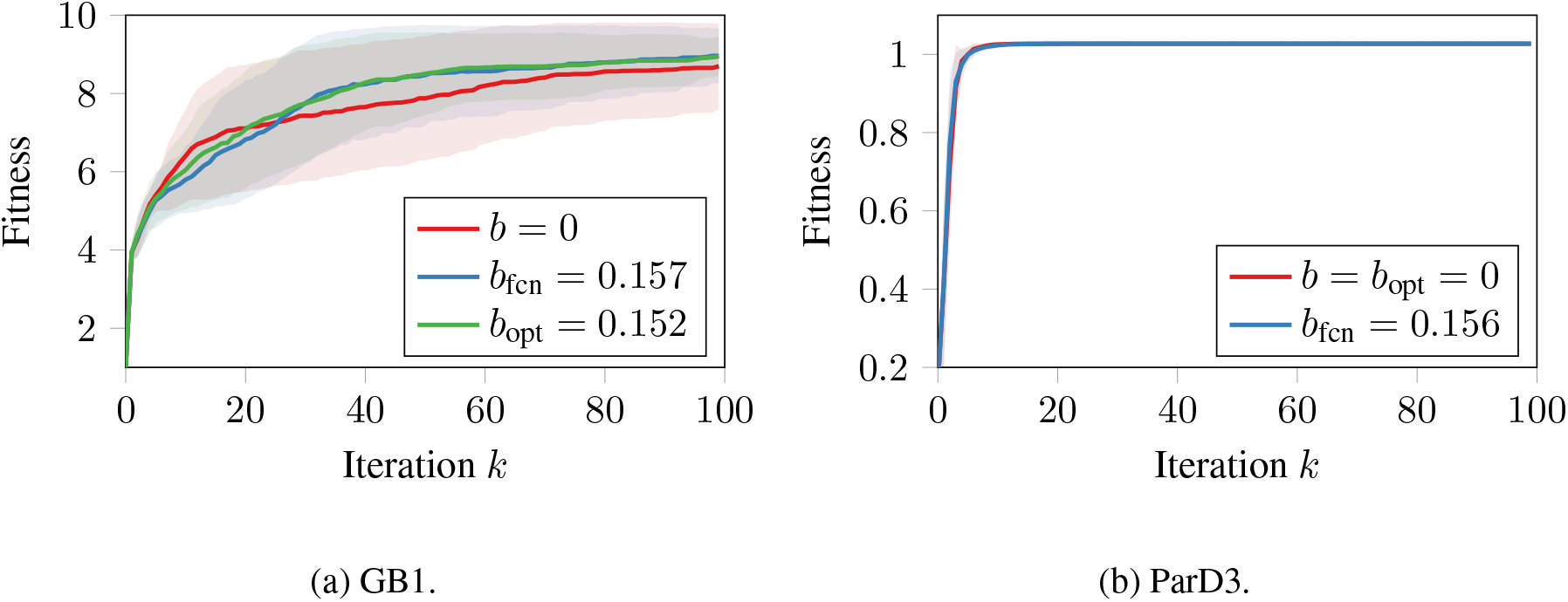
Mean fitness and standard deviation for simulated directed evolution experiments with population size *n* = 300 averaged over 100 different starting points.

## 5. Conclusion

In this paper, it was shown that the ruggedness of protein fitness landscapes can be estimated from measured average fitness values, *μ*_*k*_. The estimated ruggedness parameter can be used to select parameters for control of directed evolution. To estimate the ruggedness parameter, an FCN was trained on a range of *NK* landscapes with known ruggedness parameters. The performance of the FCN-selected parameters were tested on the GB1 and ParD3 empirical landscapes, and compared to a fixed parameter (standard) approach to directed evolution. It was shown that the proposed method leads to improvements on the more rugged GB1 landscape, and matches the already high performance of the standard approach on ParD3. In the absence of prior knowledge regarding the shape of a fitness (or other) landscape, the proposed method allows one to determine a tailored strategy that improves the likelihood of a desired outcome, contrasting fixed-parameter approaches that are prone to getting trapped in local optima.

The assumption underlying the ruggedness estimation was that *μ*_0_ significantly differs from the overall mean fitness, 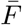, allowing the features to capture a trend as 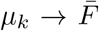. When 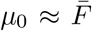, the features observed in smooth and rugged landscapes do not differ, in which case the FCN performs worse. Although in practice it is unlikely that 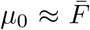 for larger *N*, future research could address this limitation, e.g., by adding higher moments of the fitness distribution to the features.

We believe that the ruggedness estimation can be improved by increasing the quality of training data, e.g., with a fitness landscape model more tailored to the case of protein evolution than the *NK* landscapes. Future research could develop a theoretical fitness landscape model that incorporates the specific properties of proteins and their evolution. This method could be validated in real-world directed evolution experiments, which commonly feature much larger values of *N* than the empirical landscapes used in this paper. Additionally, future research could combine the ruggedness estimation with the base chance look-up to obtain an end-to-end solution for parameter selection.

Although this method has been applied to the literal case of directed evolution, it could also apply to other non-linear, non-convex optimisation problems, e.g., for controlling the parameters of genetic algorithms.

## Appendix A.

In order to prove Proposition 1, several intermediary definitions and results are provided. First, the random process corresponding to a mutating gene is defined. Note that the more general continuous time setting is used in the following.

### Definition 2

**(Randomly drifting locus)** *Let* 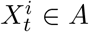 ∈ *A be a random process*. 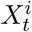 *is referred to as a randomly drifting locus with A alleles, and mutation rate α if it is a continuous-time Markov chain, with an transition rate of* 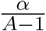 *between any two non-identical states*.

### Lemma 3

*If* 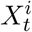 *is a randomly drifting locus with A alleles, and mutation rate α, then*, 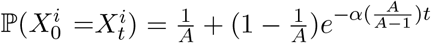.

**Proof** All alleles not equal to 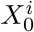 are symmetric. Hence we may consider them as a single collective state, and by symmetry the chance that 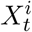 will be any specific allele given 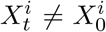 will be 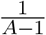 by symmetry. The result from Lemma 3 may be obtained from solving the following differential equation:

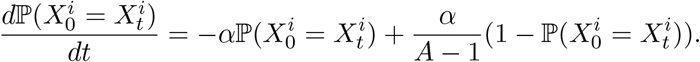

The following definition extends Definition 2 to genotype level.

### Definition 4

**(Randomly drifting gene)** *Let* 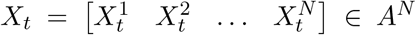 *be a random process. If all of the processes* 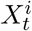 *are statistically independent, randomly drifting loci with A alleles and mutation rate α, then X*_*t*_ *is referred to as a randomly drifting gene on N loci, A alleles, and mutation rate α. This may be written shorthand as X*_*t*_ ∼ D(*N, A, α*)

### Lemma 5

*Any subset of the loci of a randomly drifting gene is also a randomly drifting gene*.

**Proof** This follows from Definition 4.

### Lemma 6

*If X*_*t*_ ∼ 𝒟 (*N, A, α*), *then* ℙ(*X*_*t*_ = *X*_0_) ≈ *e*^−*αNt*^.

**Proof** 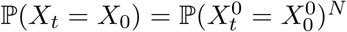, by independence of the loci. Using Lemma 3 and approximating *e*^*x*^ ≈ 1 + *x*, we obtain:

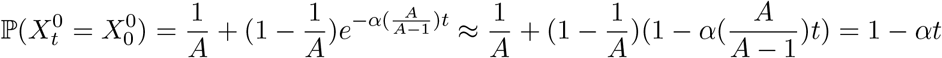

Again approximating *e*^*x*^ ≈ 1 + *x* yields the desired result for ℙ(*X*_*t*_ = *X*_0_) as

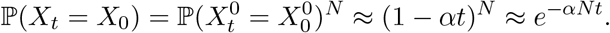

Next, the conditions that the fitness landscapes must satisfy for Proposition 1 to hold are formally defined.

### Definition 7

(*K***-loci landscape)** *We say F* : *A*^*N*^ → ℝ *is a K-loci landscape if* ∃Φ^*v*^ : *A*^*K*^ →ℝ ∀ *v*∈P_*K*_ (*N* ) *such that* ∀*X* = [*X*^1^ *X*^2^ … X^*N*^]∈ A^N^:

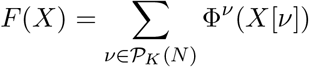

*where 𝒫*_*K*_(*N* ) = {*v* ⊆{1 … *N*{ |*K* = |*v*{, *and X*[*v*] *A*^*K*^ *is shorthand for* 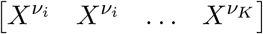, *where v*_*i*_ *is shorthand for element i of v (with the standard ordering)*.

### Definition 8

**(Isotropic** *K***-loci landscape)** *Let F* : *A*^*N*^ → ℝ *be a distribution over K-loci landscapes. We say F is isotropic if*, ∀*v*^1^, *v*^2^ ∈ 𝒫_*K*_(*N* ) *and* ∀*X*_1_, *X*_2_ ∈ *A*^*N*^ :

1. Φ^*v*1^ (*X*_1_[*v*_1_]) *and* Φ^*v*2^ (*X*_2_[*v*_2_]) *are independent unless v*_1_ = *v*_2_ *and X*_1_[*v*_1_] = *X*_2_[*v*_2_].
2. Φ^*v*1^ (*X*_1_[*v*_1_]) *and* Φ^*v*1^ (*X*_2_[*v*_1_]) *have the same distribution*.

*Furthermore, we write*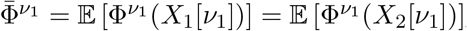, *and* 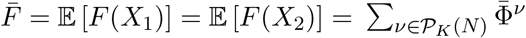

### Lemma 9

*All NK landscapes are isotropic K-loci landscapes*.

**Proof** We provide a proof by construction. Let *κ* = *κ*_1_, *κ*_2_ … *κ*_*N*_ be the *N* interaction loci in an *NK* landscape. Write 𝒰(0, 1, *k*) for the Irwin–Hall distribution, which is the sum of *k* independent standard uniform distributions. Let *k*_*v*_ = |{*i* |*κ*_*i*_ = *v*{|, for *X*∈*A*^*N*^, we may set Φ^*v*^(*X*[*v*]) ∼𝒰 (0, 1, *k*_*v*_). This induces an *NK* landscape, whilst also trivially satisfying both requirements for an isotropic *K*-loci landscape.

Finally, the main theoretical result is provided:

### Proposition 10

*Suppose X*_*t*_ ∼ D(*N, A, α*), *and F* : *A*^*N*^ → ℝ *is an Isotropic K-loci landscape. Then*, 𝔼 [*F* (*X*_*t*_) | *F* (*X*_0_)] ≈ *e*^−*αKt*^*F*_0_ + (1 − *e*^−*αKt*^) 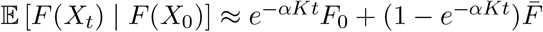.

**Proof** We write *F*_*t*_ = *F* (*X*_*t*_) as shorthand. Using Definition 8, we obtain:

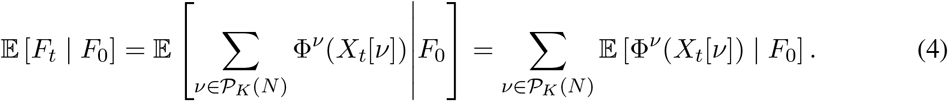

Focusing on a single *v*, by Lemma 5, *X*_*t*_[*v*] ∼ 𝒟 (*K, A, α*), allowing Lemma 6 to be used:

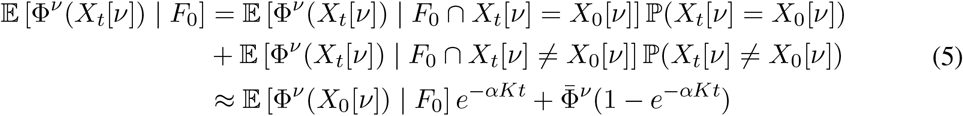

Substituting (5) into the right-hand side of (4) yields

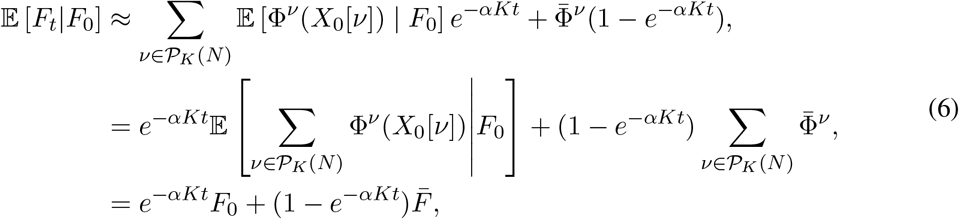

which proves Proposition 10. Proposition 1 follows from Lemma 9 and discretising (6).

## Acknowledgments

I.K. acknowledges support from EPSRC (EP/X017982/1).

## Notes

### Competing Interest Statement

The authors have declared no competing interest.

